# Enhanced soil quality after forest conversion to vegetable cropland and tea plantation has contrasting effects on soil microbial structure and functions

**DOI:** 10.1101/2021.08.07.455503

**Authors:** Lichao Fan, Guodong Shao, Yinghua Pang, Hongcui Dai, Lan Zhang, Peng Yan, Zhenhao Zou, Zheng Zhang, Jianchu Xu, Kazem Zamanian, Maxim Dorodnikov, Xin Li, Heng Gui, Wenyan Han

## Abstract

Land-use changes could potentially exert a strong influence on soil quality and soil microbial communities. Moreover, microbial taxa are also important drivers of soil ecological functions. However, the linkage between soil quality and soil microbial communities is in need of deeper understanding. In this study, we examined the effects of soil quality on microbial community structure and functions after forest conversion to vegetable cropland and tea plantations. Soil quality index was significantly increased after natural forest conversion to vegetable cropland and tea plantations. Soil bacterial beta diversity significantly correlated to soil quality, but the sensitivity of individual microbial groups varied in response to changes in soil quality. Higher soil quality promoted bacterial diversity in vegetable cropland but decreased it in tea plantations, which implied soil quality was a structural factor in bacterial community composition but had contrasting effects for croplands versus plantations. Agricultural management played a negative role in maintaining microbial interactions, as identified by the network analysis, and furthermore the analysis revealed key functions of the microbial communities. After land-use change, the abundance (e.g., level, intensity) of microbial N-cycling function increased in tea plantations but decreased in vegetable cropland. The abundance of C-cycling function featured an opposite trend. Higher level of N-fixation in tea plantations but the higher abundance of N-oxidation in vegetable cropland was demonstrated. Higher abundance of ammonia-oxidizing bacteria and ammonia-oxidizing archaea as identified by qPCR in vegetable cropland corroborated the FAPROTAX function prediction. Therefore, the key taxa of soil microbial communities and microbial functions were largely dependent on changes in soil quality and determined responses to specific agricultural management.

## 1. Introduction

Land-use change is the most impactful factor on the surface cover and soil quality of ecosystems (Foley et al., 2005). Moreover, land-use changes modify soil microbial communities, which are important drivers for maintaining soil quality and ecological functions (Jesus et al., 2009; Lehmann et al., 2020). The ecological functions of soil microorganisms play indispensable roles in carbon (C) and nitrogen (N) cycling and are closely related to plant productivity and sustainability. Thus, microorganisms and related ecological functions impact the health of soils, plants and animals (Fierer, 2017). This necessitates understanding the effects of modifications in structure and functions of microbial communities resulting from land use change and management (Fierer, 2017; Nemergut et al., 2013).

Soil quality represents the capacity of a soil to provide ecological services, and it is an important index for revealing land sustainability and productivity (Doran, 1994). Among various natural and anthropogenic factors, the type (mineral and organic) and amount of fertilization play leading roles in maintaining or altering soil quality. For example, during the conversion from natural broad-leaf forest to tea (*Camellia sinensis* L.) plantations, mineral fertilization decreased organic carbon (SOC) stock and hence negatively affected the soil quality (Fan and Han, 2018; Zhu et al., 2020). Long-term N fertilization commonly induces soil acidification (Guo et al., 2010; Kuzyakov et al., 2021, Geoderma). Changes in the aboveground vegetation cover impact soil quality because of differences in plant physiological preferences for certain nutrients as well as differing strategies for the belowground distribution of photosynthesized C (e.g., C:N ratio) and quantity of litters (Leff et al., 2012; Pausch and Kuzyakov, 2018; Sayer et al., 2011). Therefore, types of fertilization and vegetation cover are the key parameters for determining the soil quality index.

Soil quality directly and indirectly influences soil microbial communities. For example, soil pH is one of the most powerful drivers affecting microbial diversity. Lower pH inhibits microbial multiplication and growth (Malik et al., 2018; Tripathi et al., 2018; Zhalnina et al., 2015). N availability is another notable factor impacting the diversity and structure of bacterial communities (Cederlund et al., 2014; Sul et al., 2013; Wang et al., 2018). Many studies have focused on one or several specific components (e.g., SOC, pH) of soil quality to address its effects on the diversity and abundance of soil microbial taxa (Cederlund et al., 2014; Malik et al., 2018). However, studying individual soil properties may not accurately represent the holistic soil quality-related ecological functions of microorganisms (Guo et al., 2020). Soil quality is a multifactorial index that characterizes the potential of a soil to sustain its ecological services. Microbial communities, in turn, impact soil quality via nutrient cycling. But changes in microbial structure and functions may occur faster than noticeable changes in soil quality index (Bünemann et al., 2018). This is why soil quality and microbial diversity are closely correlated and interactive (Guo et al., 2020; Ji et al., 2020). However, microbial parameters are often not included in the commonly used soil quality index. In this study, we explored the links between soil quality and the complexity (diversity, structure, and functions) of microbial communities under land-use change from natural forests (unmanaged ecosystems) to vegetable cropland or tea plantation.

Tea (*Camellia sinensis* L.) is a perennial evergreen broad-leaf cash crop, and tea plantations in eastern China are typically established after conversion from forests. Tea plantations are currently covering 3.06 million ha in China alone and rapidly expanding in the world (Fan and Han, 2020). Tea plants are well adapted to grow in acidic soils (pH < 5.5) in contrast to other crops (5.5 < pH < 8.9) (Guo et al., 2010). Therefore, changes in diversity, structure and functions of soil microorganisms are expected with conversion from natural forest to tea plantations when compared to other crops such as vegetables.

We have analyzed microbial communities via pyrosequencing the soils of a natural broad-leaf forest, a vegetable cropland, and three tea plantations with high, medium, and low productivity (classified based on fertilization amounts and tea yields; see details in Materials and Methods). Our research questions are (i) how does soil quality respond to land-use change from forest to vegetable cropland and tea plantations? as well as (ii) does the structure and function of microbial communities depend on soil quality? The main goal was to explore linkages between soil quality and microbial ecological functions for improving sustainable agriculture.

## 2. Materials and Methods

### 2.1 Site description and soil collection

The experimental fields were established at the Chinese Academy of Agricultural Sciences Tea Research Institute, Hangzhou City, Zhejiang Province, Eastern China (120°09’E, 30°14’N), a region with a subtropical humid monsoon climate. The mean annual air temperature of the region is 17.0 °C, and the mean annual precipitation is 1533 mm (Fan and Han, 2020). The experimental fields included three land-use types: (i) a broad-leaf evergreen forest; (ii) a household vegetable field; and (iii) tea plantations (Table 1). The tea plantations were further divided in three groups based on fertilization amount and yield as: high tea productivity (HP_tea), medium tea productivity (MP_tea), and low tea productivity (Low_tea).

**Table 1.** Land-use types and practices of the experimental fields. For tea plantations, HP, MP and LP mean high-, medium- and low productivity, respectively.

| Land use type |  | Land use practice |  |  |  |  |  |
| --- | --- | --- | --- | --- | --- | --- | --- |
|  |  | Soil temperature (□) | Soil water content (%) | Mineral fertilizers as urea and compound fertilizers (kg N ha<sup>-1</sup> yr<sup>-1</sup>) | Organic fertilizers as farmyard manure and (biogas slurry in vegetable field, and rape see cake in tea plantations) ( kg ha<sup>-1</sup> yr<sup>-1</sup>) | Litter, residues or pruned trimmings (t ha<sup>-1</sup> yr<sup>-1</sup>) | Tea yield (ha<sup>-1</sup> yr<sup>-1</sup>) |
| **Forest** | *Schima crenata* Korthals, *Castanopsis sclerophylla* and *Cinnamomum camphora* | 24.0 | 29.6 | NA | NA | 10 | NA |
| **Vegetable field** | Chinese cabbage (*Brassica rapa chinensis*) and radish (*Raphanus sativus*) | 26.1 | 15.6 | 300 | 10 000 | 4 | NA |
| **HP_tea** | *Camellia sinensis* L. Longjing43 | 24.7 | 27.0 | 600 | 2 250 | 13.4 | 280 |
| **MP_tea** | *Camellia sinensis* L. Longjing43 | 25.0 | 26.2 | 450 | 1 120 | 9.4 | 200 |
| **LP_tea** | *Camellia sinensis* L. Longjing43 | 25.8 | 23.3 | 300 | 1 120 | 6.2 | 150 |

The soil at the experimental field is classified as Ultisols (US Taxonomy) which has been developed on Anshan quartz free porphyry parental material (Han et al., 2007). The vegetable field and tea gardens were transformed from forest approximately 35 years ago. The vegetable field was covered by plants throughout most of the year, and 1 t ha^-1^ lime was applied after deforestation to raise the pH value. In tea gardens, tea is harvested from mid-March to mid-April for the spring production of Westlake *Longjing* green tea. Afterward, heavy pruning management is applied to leave *in situ* as surface mulch. Field management including weeding and a fertilization schedule were similar in tea plantations, though fertilization amounts varied. The distance between the five experimental fields is less than 1 km. Detailed information on types of vegetation and land-use management can be found in Table 1. In each field, we set up three 20 m×20 m plots. Six soil samples were randomly collected from each plot at a depth of 0–20 cm, using an auger to form a composite. In total, 15 soil samples (5 land-use fields × 3 replicated plots) were collected. All soil samples were sieved through 2 mm in the field, transferred to laboratory within an hour and stored at −18 °C until analyses. A portion of the samples was air dried for soil properties analyses.

### 2.2 Soil physiochemical analyses

Air-dry soil samples were used to measure pH, SOC, total N content, exchangeable P and K. Moist soil samples from the field were used to determine microbial biomass, water dissolved organic C (DOC), water dissolved nitrogen (DON), NO_3_^-^-N, and NH_4_^+^-N. For details on measurement methods please refer to (Fan et al., 2015; Fan and Han, 2020). In brief, a combined glass electrode (Orion 3-Star Benchtop pH Meter; Thermo Scientific Waltham, MA, USA) was used to determine soil pH; soil total N and SOC content were determined by a Vario Max CN Analyzer (Elementar Analysensysteme GmbH, Langenselbold, Germany); a C/TN analyzer (multiN/C 2100; Analytikjena, Jena, Germany) were used to measure DOC and DON; fumigation–potassium sulfate (K_2_SO_4_) extraction method used for MBC and MBN determination where MBC and MBN were calculated from the difference between the extracted organic C and N content of fumigated and un-fumigated soils using a *k_ec_* factor of 0.45 (Joergensen, 1996). Exchangeable P was extracted with 0.03 mol L^-1^ ammonium fluoride and 0.025 mol L^-1^ hydrochloric acid; available soil K was extracted with 1 mol L^-1^ ammonium acetate; all extractions were done in a 1:10 soil-water ratio, and the oscillation was 30 min. The elements K and P were determined using inductively coupled plasma atomic emission spectroscopy (ICP-AES; JAC IRIS/AP, Thermo Jarrell Ash Corporation, Franklin, USA). NO_3_^-^-N and NH_4_^+^-N were extracted with 0.05 M CaCl_2_ and measured using continuous flow injection colorimetry (Flow Access 12.0, Skalar, Dutch). The measured soil physiochemical properties are showed in Table S1.

### 2.3 Soil quality index

Soil quality index based on management assessment framework was determined according to (Shao et al., 2020) as follows: the principal component analysis (PCA) was performed to select the minimum data set (MDS) of properties (Table S1), which could best represent soil quality and are sensitive to land management (Nakajima et al., 2015). Significant principal components (PCs) (eigenvalues ≥ 1, Table S2) were chosen to weigh properties, and a property was selected when its score value was above 10% of the highest indicator (Andrews and Carroll, 2001). Notably, the indicator with highest score value in PC was selected into the MDS when the above selected indicators were multicollinear (*p* < 0.05) based on the Pearson’s correlation (Table S3). Thereafter, the selected indicators were transformed and normalized to a value between 0.1 and 1.0 using the standard score function method. Three standard scoring functions (i) “more is better” (ii) “less is better” and (iii) “optimum” were applied to standardize the MDS indicators. The weight of indicators in MDS based on the communality of the PCA (Table S2) was calculated as the quotient of the communality divided by the sum of the communality of indicators in the MDS (Shao et al., 2020). Finally, soil quality index was calculated as follows (Doran, 1994):

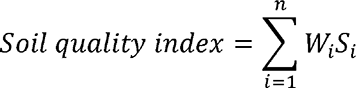

*W_i_* is the weighing value, and *S_i_* is the transformed score of the selected indicators. The quality index commonly corresponds to a proxy showing capability of a soil to produce higher yields. However, in the current study, we highlight that a higher quality index did not represent a higher capacity of soil to sustain all plant growth but intended for specific land uses (e.g., tea plantation).

### 2.4 DNA extraction and pyrosequencing

Genomic DNA was extracted from 0.5 g soil sample using the Mo Bio PowerSoil DNA isolation kit (Carlsbad, CA, USA) according to the manufacturer protocol. The DNA purity and concentration were determined using the NanoDrop ND 200 spectrophotometer (Thermo Scientific, USA). The V4-V5 variable region of the bacterial 16S rRNA gene was amplified using the primers 515F (5’-CCATCTCATCCCTGCGTGTCTCCGAC-3’) and 907R (5’-CCTATCCCCTGTGTGCCTTGGCAGTC-3’). All the polymerase chain reactions (PCR) were performed with 1 μL purified DNA template (10 ng), 5 μL 10 × PCR buffer, 2.25 mmol L^−1^ MgCl_2_, 0.8 mmol L^−1^ deoxyribonucleotide triphosphate (dNTP), 0.5 μmol L^−1^ of each primer, 2.5 U Taq DNA polymerase, and sterile filtered milli-Q water to a final volume of 50 μL. All reactions were carried out in a PTC-200 thermal cycler (MJ Research Co., New York, USA). PCR cycles included a 4 min initial denaturation at 94 □, followed by 30 cycles of denaturation at 94 □ for 1 min, annealing at 53 □ for 30 s, extension at 72 □ for 1 min, and a 5-min final elongation step at 72 □. PCR products were quality-screened and purified using the Qiangen Gel Extraction kit (Qiagen, Hilden, Germany).

The abundance of ammonia-oxidizing bacteria (AOB) and ammonia-oxidizing archaea (AOA) was determined by real-time quantitative polymerase chain reaction (qPCR) using primers that amplify *amoA* genes.

### 2.5 454 Pyrosequencing and sequencing data analysis

Pyrosequencing was performed on a Roche Genome Sequencer FLX+ using Titanium chemistry by Macrogen (Roche Applied Science, Mannheim, Germany). Three standard flow-gram format (SFF) files were generated by 454 pyrosequences. The SFF file was analyzed by the software package Mothur (version 1.33.2). The work flow was similar with (Gui et al., 2021). Briefly, De-noising was conducted with the AmpliconNoise, and UCHIME algorithms were used to reduce sequence errors and remove chimeras. Remaining sequences were aligned with the SILVA-based bacteria reference database. Sequences with 97% similarity were clustered into the same Operational Taxonomic Units (OTUs) according to the UCLUST algorithm. For each OTU, the Green Gene database was applied to annotate taxonomic information.

### 2.6 Data analyses

The least significant difference (LSD) test was used to determine differences in soil physicochemical properties between the experimental fields. The within sample alpha (α-) diversity of soil bacterial communities was calculated based on the OTU table as observed OTU number, ACE (Abundance-based Coverage Estimator metric), Chao1, and Shannon diversity index. Principal Coordinate Analysis (PCoA) based on Bray-Curtis distance between samples were calculated as the beta (β) diversity of microbial communities (compositional dissimilarity between fields) using the *ampvis2* package in R (v4.0.3). Differential OTU abundance was performed using a generalized linear model with *p* value <0.01 in the BioConductor package *EdgeR*.

The co-occurrence patterns between bacterial OTUs were explored using network analysis, and the relative abundance of OTUs above 0.05% was selected. Pairwise Pearson correlations were calculated between the remaining OTUs. A valid co-occurrence was considered as a statistically robust correlation between taxa when the Pearson’s correlation (r) was >0.6 and the *p* value was <0.01. Each node indicated individual OTU, and each edge represented the pairwise correlations between nodes standing for a significant metabolic association in the network. Multiple topological properties (i.e., number of nodes and edges, average degree) were calculated and visualized using *igraph* package in R (v4.0.3). Functions were inferred using FAPROTAX (Louca et al., 2016), which is a conservative algorithm currently matching 80 functions against 7600 functional annotations of 4600 prokaryotic taxa. Significant difference (*p* < 0.5) of functions between fields was revealed by STAMP (Statistical Analysis of Taxonomic and Functional Profiles) (Parks et al., 2014), and the multiple test correction were used *Bonferroni* method.

### 2.7 Data availability

Sequencing data are available in the NCBI SRA data repository under project No. PRJNA750877.

## 3. Results

### 3.1 Soil quality index

The PCA analysis showed that samples from forest, vegetable field, and tea gardens distributed in different quadrants featured a higher degree of soil heterogeneity after land use change and a strong difference between vegetable fields and tea plantations (Fig. S1). Indeed, soil quality index was highest in vegetable cropland with Chinese cabbage and radish (Table 1), followed by tea plantations and forest (Fig. 1). Among tea plantations, soil quality index was highest for high tea productivity (HP_tea). There was no significant difference between medium and low tea productivity plots (MP_tea vs. LP_tea). Among the measured properties, soil pH, DOC, NH_4_^-^-N, NO_3_^-^-N and available P were the significant factors for the soil quality (Table S2).

**Fig. 1.**
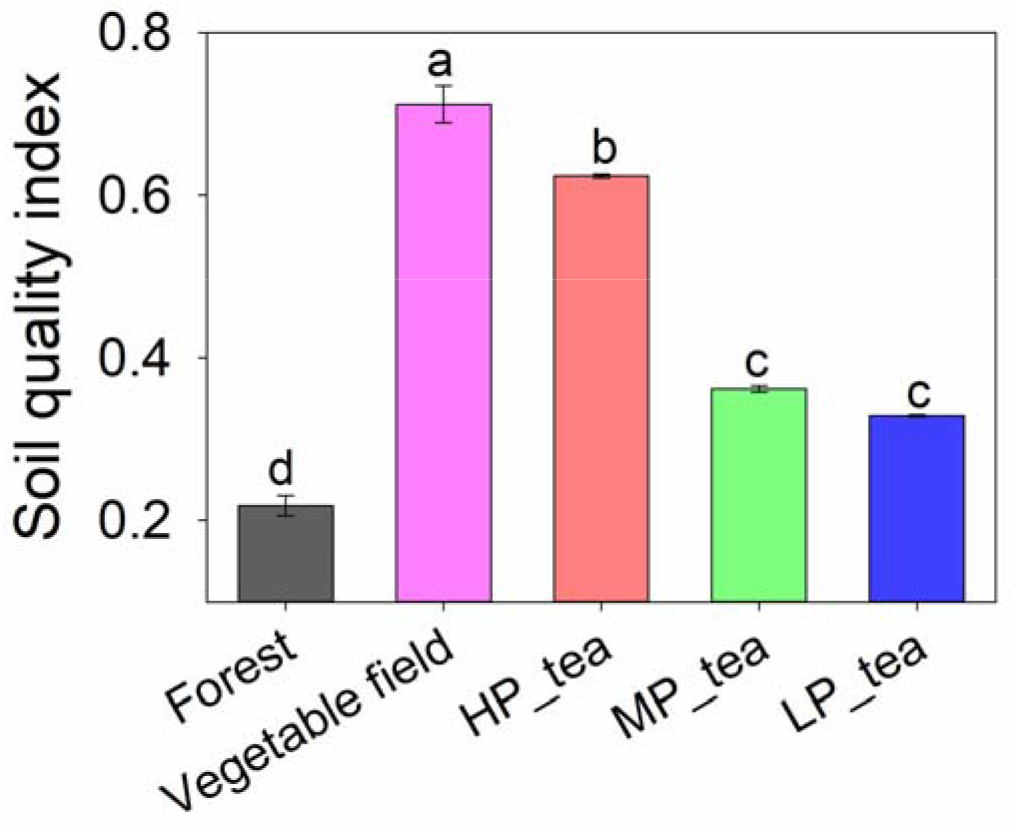
Soil quality index in forest, vegetable field, and tea plantations with high productivity (HP_tea), medium productivity (MP_tea), and low productivity (LP_tea). Lowercase letters represent significant differences at *p* < 0.05 between land uses by LSD.

### 3.2 Distribution of taxa and phylotypes across land-use types

99.6% of sequences were classified into the phyla of bacteria, and the total OTU number was 5662 defined by 97% sequence similarity. Three phyla including Proteobacteria, Acidobacteria, and Actinobacteria were predominant (relative abundance of each > 10%) (Fig. 2b). The most abundant phylum Proteobacteria, with an average relative abundance of 35.2%, included the following classes: Alphaproteobacteria (17.3%), Gammaproteobacteria (11.9%), Betaproteobacteria (3.9%), and Deltaproteobacteria (2.2%). The relative abundance of Proteobacteria, Bacteroidetes, and Nitrospira were positively correlated, but the abundance of Acidobacteria, Chloroflexi, and Armatimonadetes were negatively correlated to the soil quality index (Fig. 2a).

**Fig. 2.**
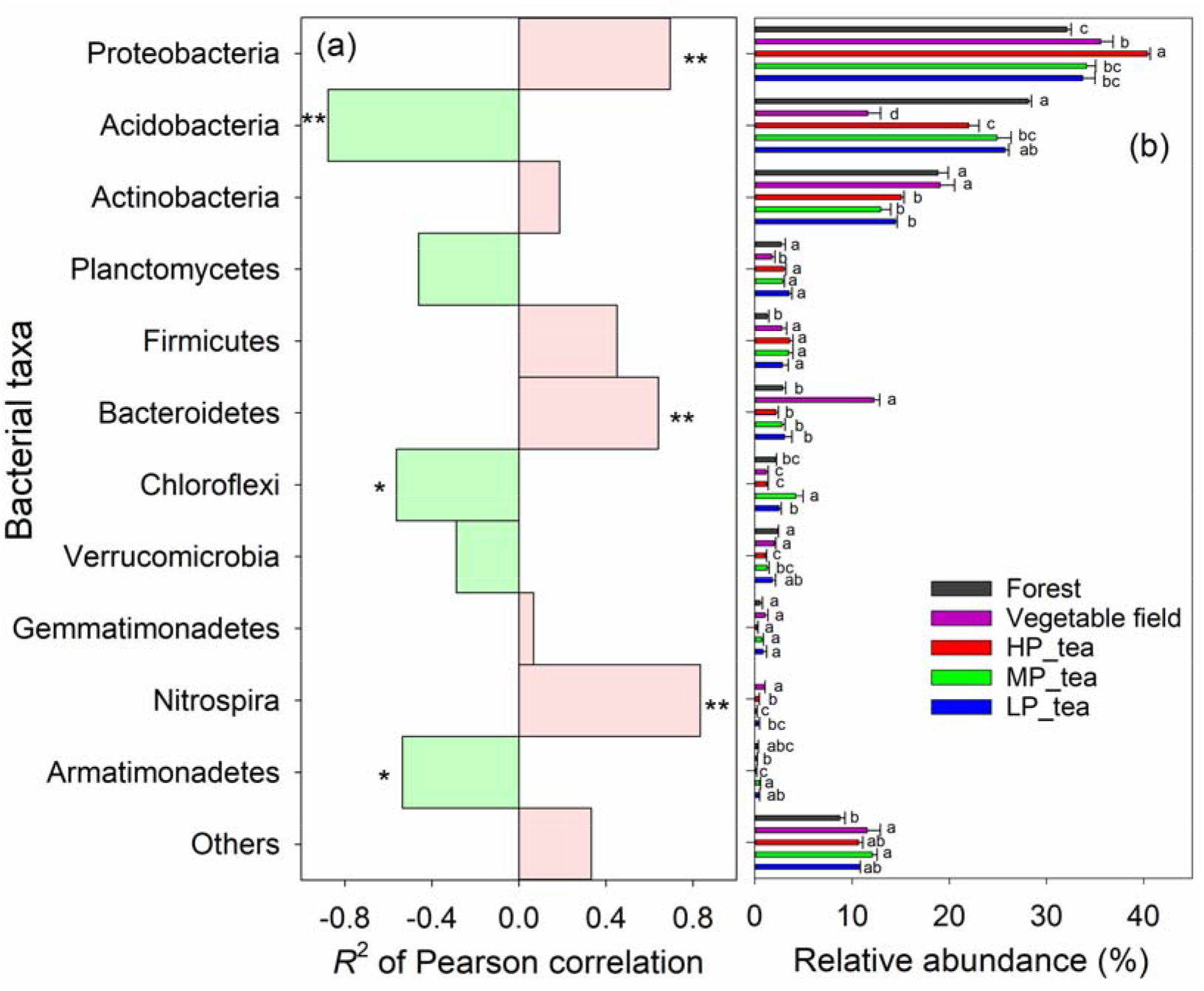
Relative abundance of top eleven phyla in forest, vegetable filed, and tea plantations with high productivity (HP_tea), medium productivity (MP_tea), and low productivity (LP_tea). Lowercase letters represent significant differences at *p*<0.05 between land uses by LSD.

Using OTU counts from forest soil as a control and an adjusted *p* value cutoff of 0.01, there were 215 OTUs enriched and 252 OTUs depleted in samples of vegetable cropland. Comparatively, the amount of enriched and depleted OTUs was much lower in tea plantation than in vegetable cropland (Fig. 3a). The number of enriched and depleted OTUs showed weak positive correlations with soil quality (Fig. S2). Noteworthy, there were overlaps in enriched OTUs between soil samples of vegetable field and tea gardens (89 out of the 215 OTUs enriched) (Fig. 3b). The enriched OTUs mainly consisted of Proteobacteria and Acidobacteria (Fig. 3d). However, 248 out of 252 OTUs were depleted only in samples of vegetable cropland (Fig. 3c), and Proteobacteria was the most depleted phylum (Fig. 3e). Therefore, land use change from forest to tea plantation had less effect on microbial communities compared with vegetable cropland.

**Fig. 3.**
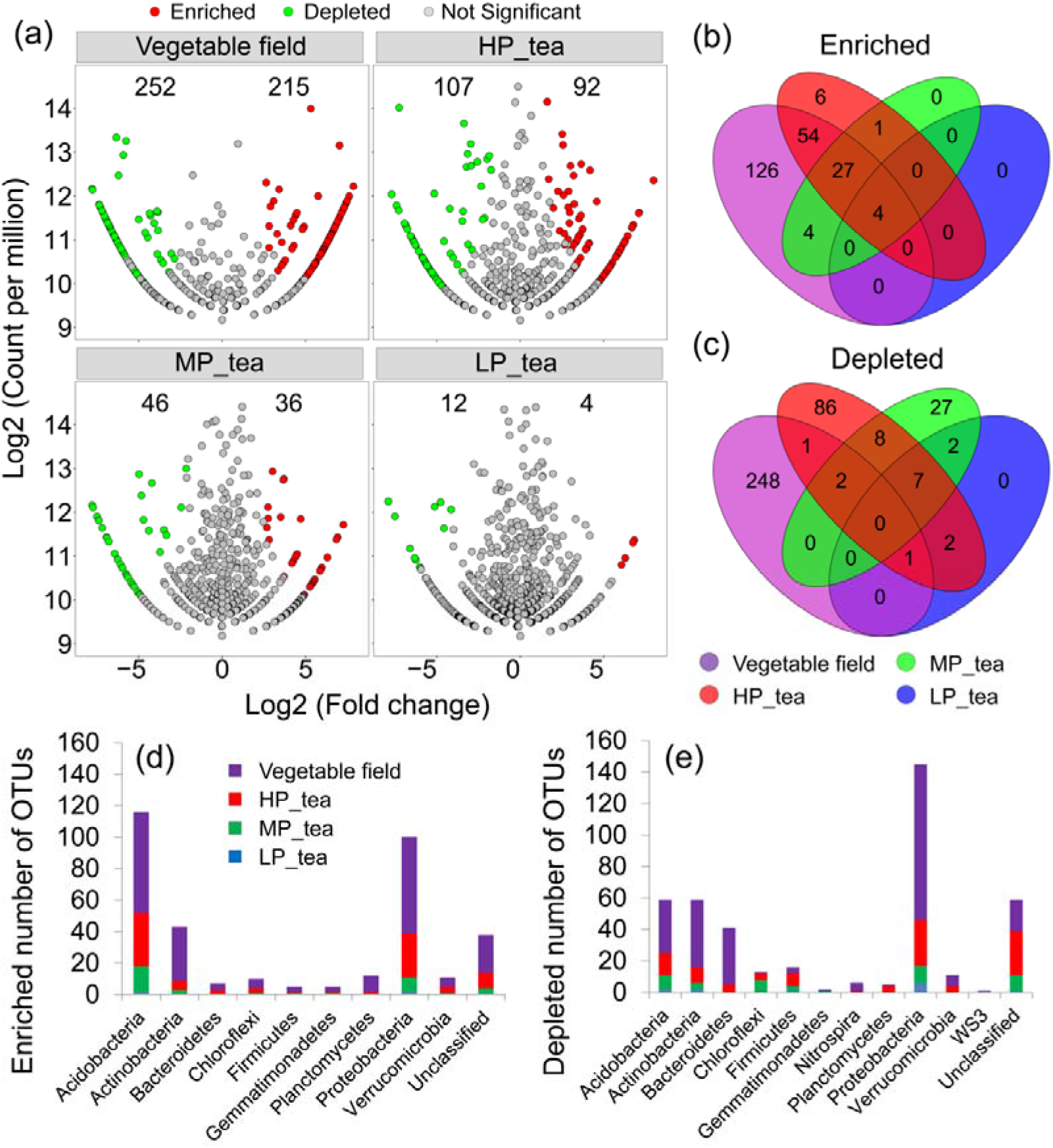
Enriched and depleted OTUs in vegetable field and tea plantations compared to forest. (a) Enrichment and depletion in fields as compared with forest. Each point represents an individual OTU, and the position along the x axis represents the abundance fold change compared with forest. (b) and (c) Shared numbers of differentially enriched and depleted OTUs between each field as compared with forest. (d) and (e) Number of depleted and enriched OTUs in each phyla.

### 3.3 Correlations between bacterial diversity and soil quality index

The α-diversity, which indicates the richness and diversity of microbial communities, was generally highest in soils of vegetable cropland than in other soils (Fig. 4a). Among tea plantations, α-diversity increased with decreasing amounts of fertilization and tea yields. We found positive correlations between soil quality and α-diversity after land use conversion from forest to vegetable field, but negative correlations after land use change from forest to tea plantations were observed (Fig. 4b). Bacterial communities were tightly associated with respective ecosystems and land use types, explaining 74.8% of bacterial community variation (PCoA analysis, Fig. 4c). 64.7% of bacterial variation among fields was significantly related to the soil quality changes (Fig. 4d). Microbial diversity was strongly dependent on the soil quality: there were distinct effects of soil quality on microbial diversity between vegetable filed and tea plantations because of different soil characteristics (Table S1). Soil pH and NH_4_^+^-N were the most important variables for explaining diversity dissimilarity in bacterial communities (RDA analysis, Fig. S3; Mantel test, Fig. S4; Pearson correlation, Fig. S5).

**Fig. 4.**
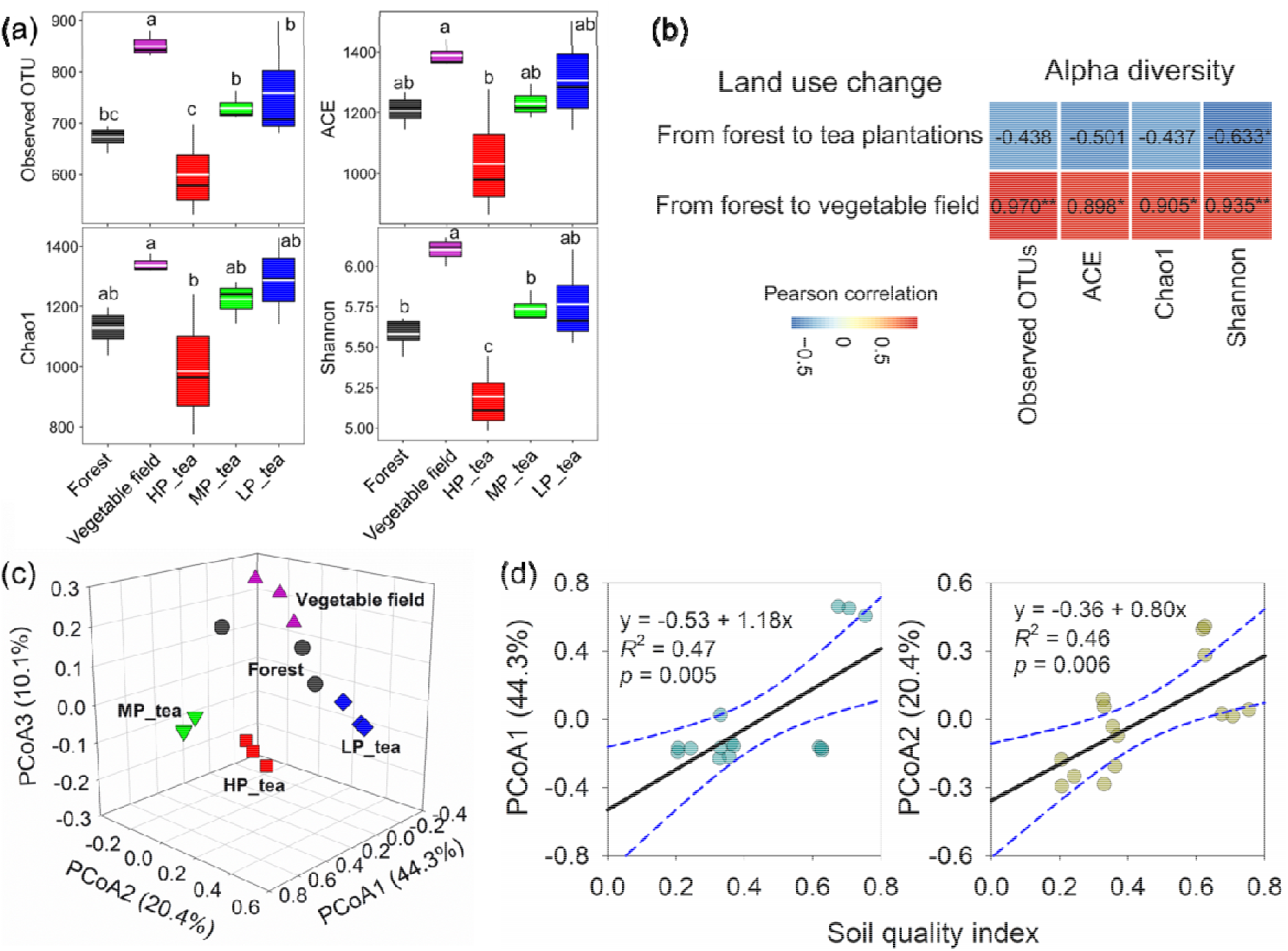
Microbial diversity in forest, vegetable filed, and tea plantations with high productivity (HP_tea), medium productivity (MP_tea), and low productivity (LP_tea). (a) Within sample alpha diversity. Lowercase letters represent significant differences at *p* < 0.05 between land uses. (b) Pearson correlations between the soil quality index and bacterial alpha diversity. ** represents the correlation is significant at the 0.01 level, * represents the correlation is significant at the 0.05 level. (c) Beta diversity. (d) Relation Blue dotted lines represent the 95% confidence interval of regression analyses.

### 3.4 Co-occurrence network analysis

The nodes in the network were assigned to eleven bacteria phyla (Fig. 5a, Fig. S6). Among these, six phyla (Acidobacteria, Proteobacteria, Actinobacteria, Verrucomicrobia, Bacteroidetes, and Firmicutes) were widely distributed, accounting for 84% of all nodes. Acidobacteria was the most abundant phyla in the network of forest soil, but Proteobacteria was dominant in vegetable field and tea plantations. The highest abundance of Nitrospira with nitrite-oxidizing bacteria responsible for nitrification was distributed in the bacterial networks of vegetable field (Fig. 5a).

**Fig. 5.**
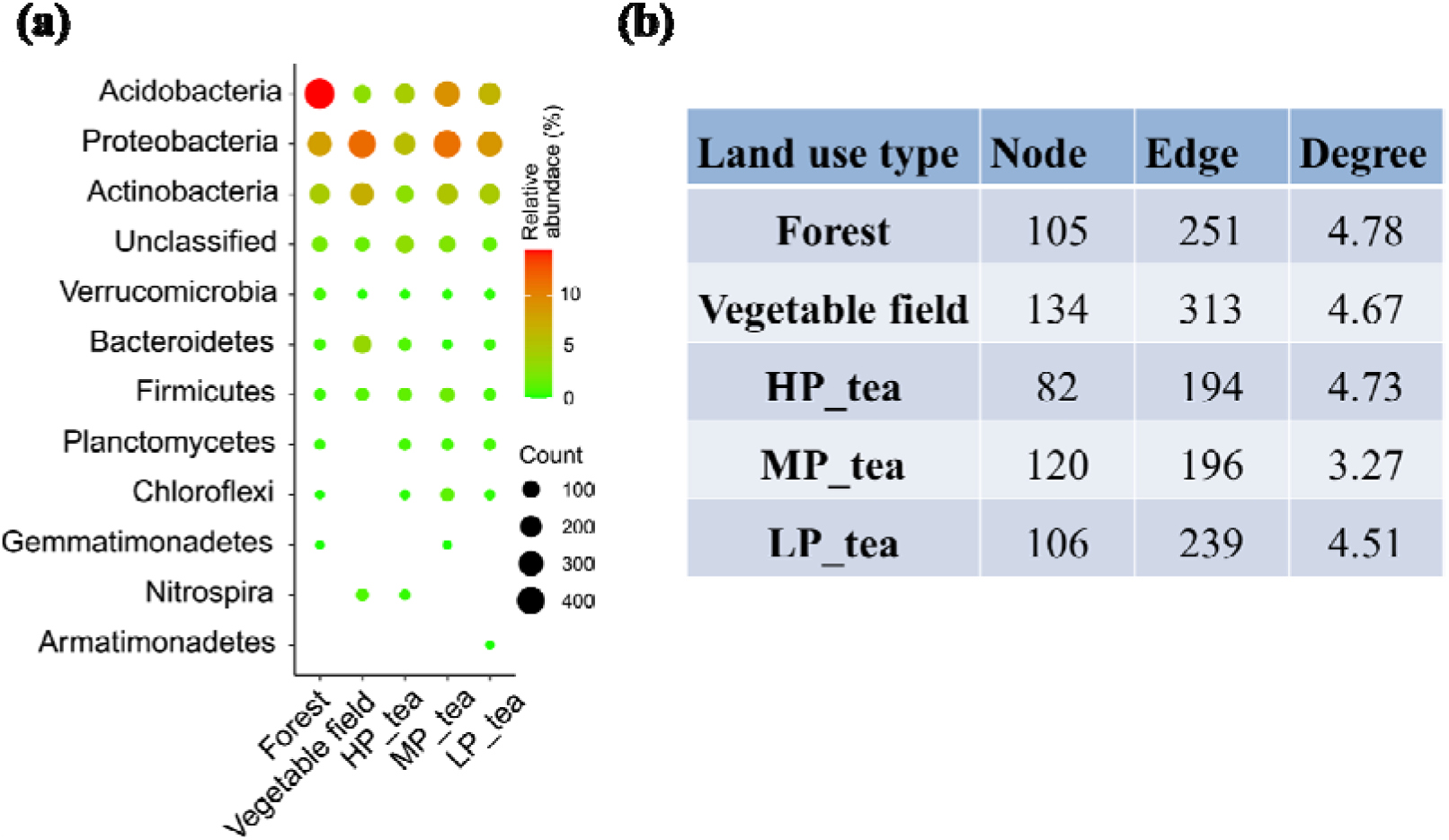
Characters of bacterial co-occurrence networks in forest, vegetable field, and tea plantations with high productivity (HP_tea), medium productivity (MP_tea), and low productivity (LP_tea). (a) The relative abundances and amounts of nodes contributed into co-occurrence networks grouped by phyla. (b) The number of nodes, edges, and degree in each network.

Compared to forest, the number of edges was larger in a network of vegetable field but lower tea plantations (Fig. S7). The degree (the number of edges per node, which implies the interactions between microbial communities) decreased in both vegetable field and tea plantations (Fig. 5b). This indicates weaker response of soil bacterial taxa to field managements (e.g., fertilization) after land use conversion from natural forest to vegetable cropland and tea plantations.

### 3.5 Microbial functions related to C- and N-cycling

Thirty-seven microbial functional categories out of 80 functions from FAPROTAX (a functional annotations dataset) were assigned. Therein, 11 microbial functions were clustered in C-cycling and 12 microbial functions were clustered in N-cycling (Fig. 6a). Seven microbial functions of C-cycling and three microbial functions of N-cycling were identified to differ significantly among fields (Fig. 6c, STAMP analysis).

**Fig. 6.**
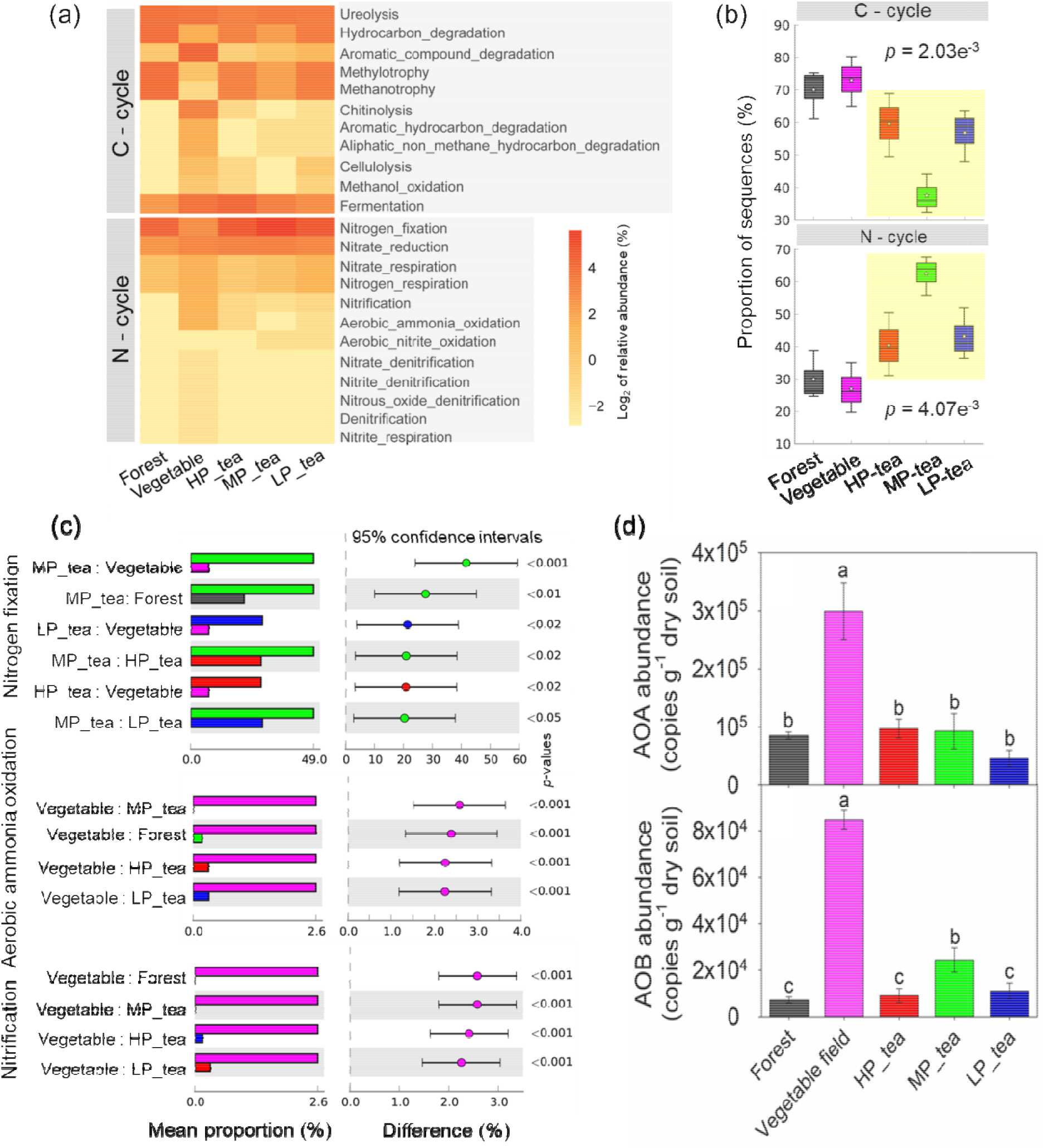
Abundance of ammonia oxidizing bacteria (AOB) and archaea (AOA) in forest, vegetable field, and tea plantations with high productivity (HP_tea), medium productivity (MP_tea), and low productivity (LP_tea). Box plots: upper and lower bars represent maximum and minimum observations, respectively; top and bottom of boxes represent third and first quartiles; thin horizontal solid lines in boxes represent median values; stars represent mean values. Lowercase letters represent significant differences at *p* < 0.05 between land uses.

The abundance of microbial C-cycling functions increased after land use change from forest to vegetable field, but decreased after land use change from forest to tea plantations. The abundance of microbial N-cycling functions was significantly higher in tea plantations than in vegetable field. Subsequently, we found that the abundance of N-fixation (i.e., Nitrogen_fixation) was higher in tea plantations than in vegetable field, but the abundance of N-oxidation (i.e., Nitrification, Aerobic_ammonia_oxidation) was higher in vegetable field than in tea plantations (Fig. 6c). Moreover, the relative abundance of ammonia-oxidizing bacteria (AOB) and ammonia-oxidizing archaea (AOA) were significantly higher in soil samples of vegetable field than in tea plantations (Fig. 6d), which corroborates the function prediction by FAPROTAX. Significant correlations between soil quality index and relative abundance of AOB and AOA were observed (Fig. S8), demonstrating the contrasting effects of land use types on microbial functioning groups.

## 4. Discussion

Soil quality index significantly increased after natural forest conversion to vegetable cropland and tea plantations (Fig. 1), directly addressing the first research question. Land management-driven changes in soil quality index were higher in vegetable cropland than tea plantations. Soil pH and NH_4_^-^-N, which were largely affected by land use types, were the most important factors in driving bacterial diversity dissimilarity and soil quality changes because pH strongly affects the availability of soil nutrients and stability of soil aggregates (Slessarev et al., 2016); moreover, pH can also directly influence the structure of microbial communities (Figs. S3-5). For example, the abundance of Acidobacteria phyla (which are particularly abundant in acidic soils) was significantly lower in vegetable cropland than the two tea plantations (Fig. 2). NH_4_^+^ is the ready source of N for microorganisms, which implies that soil microorganisms were most likely N-limited in tea plantations (see below).

Microbial communities respond rapidly to changes in soil quality caused by land use change and agricultural managements (Ji et al., 2020; Malik et al., 2018). The significant and positive correlations between the bacterial community and the soil quality index (Fig. 4d) demonstrates that soil quality is a responsible factor in assembling the microbial community structure (Guo et al., 2020), which positively supports our second research question. The linkages between soil quality and dominant bacterial taxa were either positive (i.e., Proteobacteria, Bacteroidetes, and Nitrospira) or negative (i.e., Acidobacteria, Chloroflexi, and Armatimonadetes) or insignificant (Fig. 2a), which demonstrates that individual microbial taxa respond differently to soil quality changes. Positive correlation between soil quality and bacterial α-diversity after forest conversion to vegetable field and negative correlations following conversion to tea plantations (Fig. 4b) implies that soil quality as a structuring factor in bacterial community composition has contrasting effects in croplands and plantations. Higher soil quality promoted bacterial diversity in vegetable cropland but inhibited bacterial diversity in tea plantations. This is inconsistent with fungal diversity response to soil quality. Delgado Baquerizo et al. (2017) showed that soil fungal biodiversity was positively and strongly related to soil quality at the continental scale; however, Guo et al. (2020) illustrated that the linkage between soil quality and fungal biodiversity was weaker in agricultural ecosystems than in natural ecosystems. So, linkages between microbial community and soil quality were not only affected by the types of land-use but also by the intensity of land use managements (e.g., anthropogenic pressure) (Delgado Baquerizo et al., 2017).

The interactions (i.e., degree of network) of soil bacterial communities became weaker in response to field managements (e.g., fertilization) after land use conversion from natural forest to vegetable field and tea plantations (Fig. 5), which demonstrates that land use management plays a negative role in maintaining microbial interactions, since mineral and organic fertilizers provide large amounts of ready-to-use nutrients or easily decomposed substrates, which may reduce the mutualistic network of microorganisms (Berry and Widder, 2014).

Previous studies have shown that C and N availability can influence the structure of soil bacterial communities (Cederlund et al., 2014; Sul et al., 2013; Wang et al., 2018) because soil microorganisms are generally N limited or co-limited by C and N sources (Chen et al., 2018; Wardle, 1992). Increasing microbial C-cycling in vegetable field in contrast to tea plantation (Fig. 6b) indicates that microbial functional groups developed directionally and deterministically due to the specific land managements, which drives variations in soil quality under different land uses and changes in microbial function groups. Higher abundance of N-fixation (i.e., Nitrogen_fixation) in tea plantations and higher abundance of N-oxidation (i.e., Nitrification, Aerobic_ammonia_oxidation) in vegetable cropland (Fig. 6c) suggests contrasting N demand depends on the land-use type. Comparably, soil microorganisms were more N-limited in tea plantations than that in vegetable field. Higher abundance of ammonia-oxidizing bacteria (AOB) and ammonia-oxidizing archaea (AOA) in vegetable cropland soil measured by qPCR corroborated the function prediction by FAPROTAX. Moreover, the significant correlation between soil quality index versus the abundance of AOB and AOA (Fig. S8) demonstrates that the key taxa of microbial communities were strongly dependent on soil quality. Land-use changes drive changes in soil quality, depending on management, which reshapes the microbial community. Changes in soil microbial communities in turn control various soil biogeochemical cycles (Kuypers et al., 2018). Accordingly, microbial parameters as sensitive factors in different land uses can be included in the commonly used soil quality index.

## 5. Conclusions

We confirmed that the soil quality index was significantly increased after natural forest conversion to agricultural land use (e.g., vegetable cropland, tea plantations). The structure and functions of microbial communities were influenced by soil quality, but individual microbial taxa respond differently to changes in soil quality. Microbial function N cycling thrives in tea plantations, whereas C-cycling thrives in vegetable croplands. Co-occurrence network analysis demonstrated agricultural management played a negative role in maintaining microbial interactions. Consequently, deciphering the relationships between microbial parameters-related soil quality and ecosystem functions provides a new opportunity for improving sustainable agriculture.

## Acknowledgements

This research was supported by the National Key R&D Program of China (2017YFE0107500). Heng Gui would like to thank the support from Yunnan Fundamental Research Projects (grant NO. 2019FB063) and the National Natural Science Foundation of China (NSFC Grant 32001296). Kazem Zamanian would like to thank the Research Fund for International Young Scientists of National Natural Science Foundation of China to K.Z. (Grant number: 42050410320). Funding for study abroad program by the government of Shandong Province for financial support for Dr. Hongcui Dai

## Notes

### Competing Interest Statement

The authors have declared no competing interest.

## References

Andrews, S.S., Carroll, C.R., 2001. Designing a soil quality assessment tool for sustainable agroecosystem management. Ecological Applications 11, 1573–1585.

Berry, D., Widder, S., 2014. Deciphering microbial interactions and detecting keystone species with co-occurrence networks. Frontiers in Microbiology 5, 219.

Bünemann, E.K., Bongiorno, G., Bai, Z., Creamer, R.E., De Deyn, G., de Goede, R., Fleskens, L., Geissen, V., Kuyper, T.W., Mäder, P., 2018. Soil quality–A critical review. Soil Biology and Biochemistry 120, 105–125.

Cederlund, H., Wessén, E., Enwall, K., Jones, C.M., Juhanson, J., Pell, M., Philippot, L., Hallin, S., 2014. Soil carbon quality and nitrogen fertilization structure bacterial communities with predictable responses of major bacterial phyla. Applied Soil Ecology 84, 62–68.

Chen, H., Li, D., Zhao, J., Zhang, W., Xiao, K., Wang, K., 2018. Nitrogen addition aggravates microbial carbon limitation: Evidence from ecoenzymatic stoichiometry. Geoderma 329, 61–64.

Delgado Baquerizo, M., Powell, J.R., Hamonts, K., Reith, F., Mele, P., Brown, M.V., Dennis, P.G., Ferrari, B.C., Fitzgerald, A., Young, A., 2017. Circular linkages between soil biodiversity, fertility and plant productivity are limited to topsoil at the continental scale. New Phytologist 215, 1186–1196.

Doran, J.W., 1994, Defining soil quality for sustainable environment. Wisconsin, US: Soil Science Society of America.

Fan, L., Han, W., 2020. Soil respiration after forest conversion to tea gardens: A chronosequence study. Catena 190, 104532.

Fan, L., Yang, M., Han, W., 2015. Soil respiration under different land uses in Eastern China. PLoS One 10, e124198.

Fan, L.C., Han, W.Y., 2018. Soil respiration in Chinese tea gardens: autotrophic and heterotrophic respiration. European Journal of Soil Science 69, 675–684.

Fierer, N., 2017. Embracing the unknown: disentangling the complexities of the soil microbiome. Nature Reviews Microbiology 15, 579–590.

Foley, J., DeFries, R., Asner, G., Barford, C., Bonan, G., Carpenter, S., Chapin, F., Coe, M., Daily, G., Gibbs, H., Helkowski, J., Holloway, T., Howard, E., Kucharik, C., Monfreda, C., Patz, J., Prentice, I., Ramankutty, N., Snyder, P., AF Foley, J., DeFries, R., Asner, G., Barford, C., Bonan, G., Carpenter, S., Chapin, F., Coe, M., Daily, G., Gibbs, H., Helkowski, J., Holloway, T., Howard, E., Kucharik, C., Monfreda, C., Patz, J., Prentice, I., Ramankutty, N., Snyder, P., 2005. Global consequences of land use. Science 309, 570–574.

Gui, H., Fan, L., Wang, D., Yan, P., Li, X., Zhang, L., Han, W., 2021. Organic management practices shape the structure and associations of soil bacterial communities in tea plantations. Applied Soil Ecology, 103975.

Guo, J., Ling, N., Chen, Z., Xue, C., Li, L., Liu, L., Gao, L., Wang, M., Ruan, J., Guo, S., Vandenkoornhuyse, P., Shen, Q., 2020. Soil fungal assemblage complexity is dependent on soil fertility and dominated by deterministic processes. New Phytologist 226, 232–243.

Guo, J.H., Liu, X.J., Zhang, Y., Shen, J.L., Han, W.X., Zhang, W.F., Christie, P., Goulding, K.W.T., Vitousek, P.M., Zhang, F.S., 2010. Significant Acidification in Major Chinese Croplands. Science 327, 1008–1010.

Han, W., Kemmitt, S.J., Brookes, P.C., 2007. Soil microbial biomass and activity in Chinese tea gardens of varying stand age and productivity. Soil Biology and Biochemistry 39, 1468–1478.

Jesus, E.D.C., Marsh, T.L., Tiedje, J.M., de S Moreira, F.M., 2009. Changes in land use alter the structure of bacterial communities in Western Amazon soils. The ISME journal 3, 1004–1011.

Ji, L., Ni, K., Wu, Z., Zhang, J., Yi, X., Yang, X., Ling, N., You, Z., Guo, S., Ruan, J., 2020. Effect of organic substitution rates on soil quality and fungal community composition in a tea plantation with long-term fertilization. Biology and Fertility of Soils 56, 633–646.

Ji, L., Yang, Y., Yang, L., Zhang, D., 2020. Effect of land uses on soil microbial community structures among different soil depths in northeastern China. European Journal of Soil Biology 99, 103205.

Joergensen, R.G., 1996. The fumigation-extraction method to estimate soil microbial biomass: Calibration of the k_EC_ value. Soil Biology and Biochemistry 28, 25–31.

Kuypers, M.M.M., Marchant, H.K., Kartal, B., 2018. The microbial nitrogen-cycling network. Nature Reviews Microbiology 16, 263–276.

Leff, J.W., Wieder, W.R., Taylor, P.G., Townsend, A.R., Nemergut, D.R., Grandy, A.S., Cleveland, C.C., 2012. Experimental litterfall manipulation drives large and rapid changes in soil carbon cycling in a wet tropical forest. Global Change Biology 18, 2969–2979.

Lehmann, J., Bossio, D.A., Kögel-Knabner, I., Rillig, M.C., 2020. The concept and future prospects of soil health. Nature Reviews Earth & Environment 1, 544–553.

Louca, S., Parfrey, L.W., Doebeli, M., 2016. Decoupling function and taxonomy in the global ocean microbiome. Science 353, 1272–1277.

Malik, A.A., Puissant, J., Buckeridge, K.M., Goodall, T., Jehmlich, N., Chowdhury, S., Gweon, H.S., Peyton, J.M., Mason, K.E., van Agtmaal, M., Blaud, A., Clark, I.M., Whitaker, J., Pywell, R.F., Ostle, N., Gleixner, G., Griffiths, R.I., 2018. Land use driven change in soil pH affects microbial carbon cycling processes. Nature Communications 9.

Nakajima, T., Lal, R., Jiang, S., 2015. Soil quality index of a crosby silt loam in central Ohio. Soil and Tillage Research 146, 323–328.

Nemergut, D.R., Schmidt, S.K., Fukami, T., O’Neill, S.P., Bilinski, T.M., Stanish, L.F., Knelman, J.E., Darcy, J.L., Lynch, R.C., Wickey, P., Ferrenberg, S., 2013. Patterns and Processes of Microbial Community Assembly. Microbiology and Molecular Biology Reviews 77, 342–356.

Parks, D.H., Tyson, G.W., Hugenholtz, P., Beiko, R.G., 2014. STAMP: statistical analysis of taxonomic and functional profiles. Bioinformatics 30, 3123–3124.

Pausch, J., Kuzyakov, Y., 2018. Carbon input by roots into the soil: Quantification of rhizodeposition from root to ecosystem scale. Global Change Biology 24, 1–12.

Sayer, E.J., Heard, M.S., Grant, H.K., Marthews, T.R., Tanner, E.V., 2011. Soil carbon release enhanced by increased tropical forest litterfall. Nature Climate Change 1, 304–307.

Shao, G., Ai, J., Sun, Q., Hou, L., Dong, Y., 2020. Soil quality assessment under different forest types in the Mount Tai, central Eastern China. Ecological Indicators 115, 106439.

Slessarev, E.W., Lin, Y., Bingham, N.L., Johnson, J.E., Dai, Y., Schimel, J.P., Chadwick, O.A., 2016. Water balance creates a threshold in soil pH at the global scale. Nature 540, 567–569.

Sul, W.J., Asuming-Brempong, S., Wang, Q., Tourlousse, D.M., Penton, C.R., Deng, Y., Rodrigues, J.L.M., Adiku, S.G.K., Jones, J.W., Zhou, J., Cole, J.R., Tiedje, J.M., 2013. Tropical agricultural land management influences on soil microbial communities through its effect on soil organic carbon. Soil Biology and Biochemistry 65, 33–38.

Tripathi, B.M., Stegen, J.C., Kim, M., Dong, K., Adams, J.M., Lee, Y.K., 2018. Soil pH mediates the balance between stochastic and deterministic assembly of bacteria. The ISME Journal 12, 1072–1083.

Wang, H., Liu, S., Zhang, X., Mao, Q., Li, X., You, Y., Wang, J., Zheng, M., Zhang, W., Lu, X., Mo, J., 2018. Nitrogen addition reduces soil bacterial richness, while phosphorus addition alters community composition in an old-growth N-rich tropical forest in southern China. Soil Biology and Biochemistry 127, 22–30.

Wardle, D.A., 1992. A comparative assessment of factors which influence microbial biomass carbon and nitrogen levels in soil. Biological Reviews 67, 321–358.

Zhalnina, K., Dias, R., de Quadros, P.D., Davis-Richardson, A., Camargo, F.A., Clark, I.M., McGrath, S.P., Hirsch, P.R., Triplett, E.W., 2015. Soil pH determines microbial diversity and composition in the park grass experiment. Microbial Ecology 69, 395–406.

Zhu, X., Jackson, R.D., DeLucia, E.H., Tiedje, J.M., Liang, C., 2020. The soil microbial carbon pump: From conceptual insights to empirical assessments. Global Change Biology 26, 6032–6039.

